# Transcriptomic Suppression of Immune and Regenerative Signalling in Skeletal Muscle of Patients with Chronic Kidney Disease

**DOI:** 10.1101/2025.07.11.664285

**Authors:** Luke A Baker, Nicholas Eastley, Robert U Ashford, Matthew Denniff, Matthew Graham-Brown, Emma L Watson

**Affiliations:** Department of Respiratory Sciences, College of Life Sciences, University of Leicester, Leicester, UK; NIHR Leicester Biomedical Research Centre (BRC), University of Leicester, Leicester, UK; Leicester Orthopaedics, University Hospitals of Leicester, UK; Department of Genetics, Genomics and Cancer Sciences, College of Life Sciences, University of Leicester, UK; NIHR Leicester Biomedical Research Centre, Leicester, UK; Department of Cardiovascular Sciences, University of Leicester, Leicester, UK

**Keywords:** Chronic Kidney Disease, Sarcopenia, Skeletal muscle, Transcriptomics, immune cell signalling, regeneration

## Abstract

**Background:** Chronic kidney disease (CKD) is a growing public health emergency with a global prevalence of approximately 14%. Sarcopenia is a common complication of CKD contributing to functional decline and poor outcomes. However, the molecular mechanisms driving muscle wasting in CKD remain incompletely understood. This study aimed to characterise the transcriptomic profile in individuals with CKD compared to healthy control counterparts, to identify key pathways implicated in muscle dysfunction.

**Methods:** Vastus lateralis muscle biopsy samples were obtained from n=10 people with CKD and n=9 healthy controls matched for age, sex, ethnicity and physical activity. Bulk RNA sequencing was performed on all samples. Differential gene expression was assessed using DESeq2 and pathway enrichments analyses were conducted using Gene Ontology (GO) and KEGG databases.

**Results:** A total of 76 genes were differentially expressed in CKD muscle (FDR < 0.05, |log_2_FC| ≥ 1), with 62 downregulated and 14 upregulated. Transcriptomic analysis revealed suppression of immune-related pathways, including leukocyte chemotaxis and macrophage-associated signalling (e.g., CD163, CXCL14, MPEG1). GO and KEGG analyses further supported downregulation of immune surveillance and inflammatory pathways. Several genes implicated in muscle regeneration (e.g., MEGF10, PODN, SOX4) were also differentially expressed, suggesting impaired regenerative signalling. Classical markers of myogenesis and protein degradation were unchanged, indicating a blunted rather than overtly inflammatory or catabolic muscle environment.

**Conclusions:** Skeletal muscle in CKD exhibits a distinct transcriptional profile marked by suppression of immune and regenerative processes. These findings refine our understanding of CKD-associated sarcopenia and may inform the development of targeted therapeutic strategies beyond conventional exercise-based interventions.

## Introduction

Chronic kidney disease (CKD) is common, affecting around 14% of adults in England ^1^, for which there is no cure. A loss of muscle mass and strength (sarcopenia, which is now recognised as a disease in its own right^2^), is a frequent complication of CKD affecting around 28% of people with more advanced disease ^3,4^. However, this starts early in the disease process^3^ and results in a downward spiral of further muscle wasting, disuse, reduced quality of life and poor outcome^5^. Importantly, skeletal muscle is highly adaptive, easily remodelled through interventions such as exercise and nutrition, thus the sarcopenia that is observed in these patients is likely to be reversible or preventable. Any strategies that would be capable of improving muscle mass are an attractive means to improve quality of life, clinical outcomes and reduce healthcare costs. Unfortunately, treatments for sarcopenia in this patient group and others are lacking, which is primarily hampered by our lack of complete understanding of the processes that underlie this condition. Currently the most effective strategy to reduce or prevent sarcopenia is exercise^6^. Whilst effective, activity levels of CKD patients are low^7^ and not all patients can or want to take part in exercise programmes. This is further compounded by the lack of formalised rehabilitation programmes available to CKD patients and therefore a lack of support exists to help patients to take part in regular exercise. This means we need to look to alternative strategies to help reduce/prevent sarcopenia in those patients not taking part in exercise. In order to design these alternative therapies for sarcopenia we must first understand in more detail the processes by which sarcopenia is initiated.

Multiple pathways and factors have been implicated in driving the loss of muscle mass in individuals with chronic kidney disease (CKD), including inflammation, oxidative stress, metabolic acidosis, insulin resistance, aberrant microRNA expression, and physical inactivity^8-12^. Microarray-based studies have previously examined skeletal muscle gene expression in this population and reported an enrichment of genes involved in inflammatory pathways^12^. However, high-throughput RNA sequencing (RNA-seq) exploring differential gene expression and underlying molecular pathways linked to sarcopenia in CKD remain limited. To address this, we performed RNA-seq on vastus lateralis muscle biopsies from 10 patients with CKD and 9 healthy controls to identify candidate pathways associated with muscle wasting.

## Methods

### Participants and recruitment

CKD patients were recruited from nephrology outpatient clinicals at Leicester General Hospital, UK between December 2013 and January 2019, as part of the ExTra CKD trial (Ref/10/H0406/50). Healthy controls were recruited from the University of Leicester and University Hospitals of Leicester NHS Trust during hospital attendance for routine planned orthopaedic surgery as part of the Explore CKD trial (15/EM/0467). Exclusion criteria included: age <18 years, physical impairment sufficient to prevent undertaking exercise or exercise testing, recent myocardial infarction, active cancer, unstable chronic conditions, HbA1C<9% and an inability to give informed consent. Approval for all studies was granted from the UK National Research Ethics Committee. All participants gave written informed written consent and trials were conducted in accordance with the Declaration of Helsinki. The samples were accessed for research purposes between September 2019 and December 2023.

### Muscle biopsy sampling and processing

Biopsies were collected from the vastus lateralis muscle under fasted conditions. CKD participants donated a biopsy using the micro biopsy technique as previously described^13^. Healthy controls had a biopsy taken at the same time of a scheduled procedure for the removal of benign intramuscular lipomas^14^ using the open biopsy technique. Following dissection of any visible fat, samples were immediately placed in liquid nitrogen and stored until subsequent analysis. Individual stored CKD samples were paired with a control sample matched for age (within two years), gender, ethnicity and physical activity levels assessed using the GP Physical Activity Questionnaire (GPPAQ).

### RNA extraction and transcriptome sequencing

RNA was extracted from 5mg tissue using the miRNeasy micro kit according to the manufacturer’s instructions (Qiagen Inc, Valencia, CA). RNA quality was assessed using the Ailgent 5400 fragment analyzer system. All library preparation and subsequent RNA sequencing was performed by Novogene (Beijing, China) using the Illumina Novoseq 6000 sequencing system using a 150bp paired end strategy. HISAT2 was used to align the sequencing reads to the reference genome (Homo sapiens GRCG38/hg38).

### Statistical analysis

The abundance of each gene was quantified using feature counts and then normalised to the total number of reads for each gene. The data was filtered and reads with adapter contamination, reads when it was uncertain nucleotides constitute more than 10% of either read, and reads with low quality nucleotides that constitute of more than 10% of either read were removed. The reads were aligned to a reference genome (Homo Sapiens(GRCh38/hg38)) using HIAT2. To determine reliability of the analysis, correlation of the gene expression levels between samples was determined by Perasons correlation. Principal component analysis (PCA) was used to evaluate intergroup differences and intragroup sample duplication. Due to the presence of biological replicates differential gene expression (DEG) analysis was performed using the DESeq2 package in R using the negative binomial distribution model. Hierarchical clustering analysis (HCA) was conducted to cluster differentially expressed genes (DEGs) and visualize their expression profiles pre and post exercise in the different groups. Sequencing depth and RNA quality were added into the analysis as covariates to control for potential confounding effects. Genes with adjusted P value <0.05 and with a |log2(FoldChange)|>=1 and a false discovery rate (FDR) of 0.05 ^15^ (using the Benjamini-Hochberg procedure) were considered differentially expressed. A hierarchical clustering analysis of DEGs was performed using the ggplot2 package in R. Functional enrichment analyses including Gene Ontology (GO) and Kyoto Encylopedia of Genes and Genomes (KEGG) pathway analysis were performed using the clusterProfiler software to determine which DEGs were significantly enriched in which GO terms or metabolic pathways. GO terms and KEGG pathways with padj < 0.05 were deemed significantly enriched.

## Results

### Participant characteristics

Participant characteristics are shown in Table 1.

**Table 1.** Participant characteristics

|  | **CKD group (n= 10)** | **Healthy control group (n=9)** |
| --- | --- | --- |
| **Number of Males (%)** | 3 (30%) | 3 (30%) |
| **Age (y)** | 63 (27-78) | 57 (29-75) |
| **eGFR (mL/min/1.73m<sup>2</sup>)** | 24 (15-32) | 86 (100-62) |
| **Haemoglobin (g/dL)** | 117 (102-161) | 137 (114-154) |
| **Albumin (g/L)** | 41 (23-47) | 38 (36-41) |
| **Comorbidities (n)** |  |  |
| **Type II Diabetes** | 0 | 0 |
| **Essential hypertension** | 3 | 0 |
| **Ischaemic Heart Disease** | 0 | 0 |
| **Valvular heart disease** | 1 | 0 |

Data are shown as median and range. Abbreviations: BMI, Body Mass Index; eGFR, CKD, chronic kidney disease; estimated glomerular filtration rate.

### RNA-seq quality assessment and sample clustering

RNA sequencing data were obtained from vastus lateralis muscle biopsies of 10 individuals with CKD and 9 healthy controls. After quality filtering and trimming, an average of ∼25 million clean paired-end reads per sample were retained (range: 22.7–28.9 million), with a mean alignment rate of over 94% to the reference human genome. The sequencing quality was consistently high across all samples, with Q30 scores exceeding 94%, GC content around 51%, and error rates between 0.02% and 0.03%. These metrics indicate excellent sequencing depth and accuracy suitable for downstream analyses.

PCA was performed to assess sample-level variability and clustering based on global gene expression profiles. PC1 and PC2 accounted for 10.37% and 8.86% of the total variance, respectively (Figure 1A). Despite the modest proportion of variance explained by these axes, samples clustered distinctly according to disease status, with CKD and control groups clearly separated along PC1. This suggests that CKD status exerts a consistent effect on the skeletal muscle transcriptome. Pearson correlation analysis of transcriptome-wide expression values showed high intra-group consistency (mean r = 0.938 for CKD and r = 0.930 for controls), with slightly lower but comparable inter-group correlation (r = 0.931), suggesting subtle but systematic differences that align with the PCA-based clustering.

**Figure 1.**
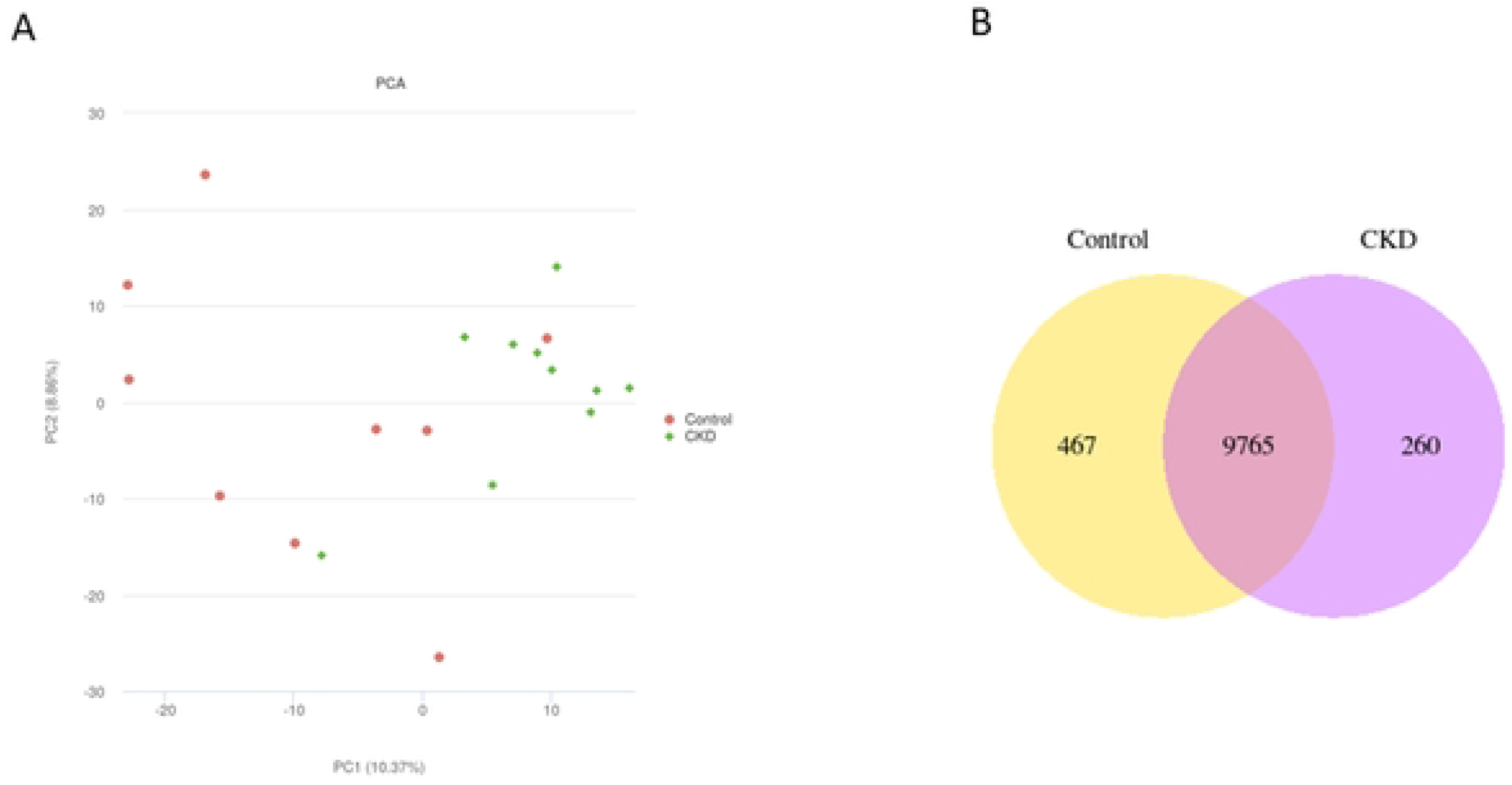
Overview of transcriptomic differences between CKD and healthy control skeletal muscle. (A) PCA plot displaying the first two principal components (PC1 and PC2) of gene expression profiles from vastus lateralis muscle biopsies of 10 individuals with chronic kidney disease (CKD; green diamonds) and 9 healthy controls (red circles). (B) Venn diagram showing the number of genes expressed (TPM > 1) in CKD and control samples.

### Gene signature of skeletal muscle from patients with CKD

On average 10,025 genes were found to be expressed (FPKM>1) in the CKD group and 10,232 genes in the healthy control group. A total of 9765 genes were shared between groups. Of these, 467 genes were uniquely expressed in the control group and 260 were unique to the CKD group (Figure 1B). Differential gene expression analysis identified a total of 76 genes that were significantly altered in CKD muscle compared to controls (FDR < 0.05), with 14 genes upregulated and 62 downregulated (Figure 2A).

**Figure 2.**
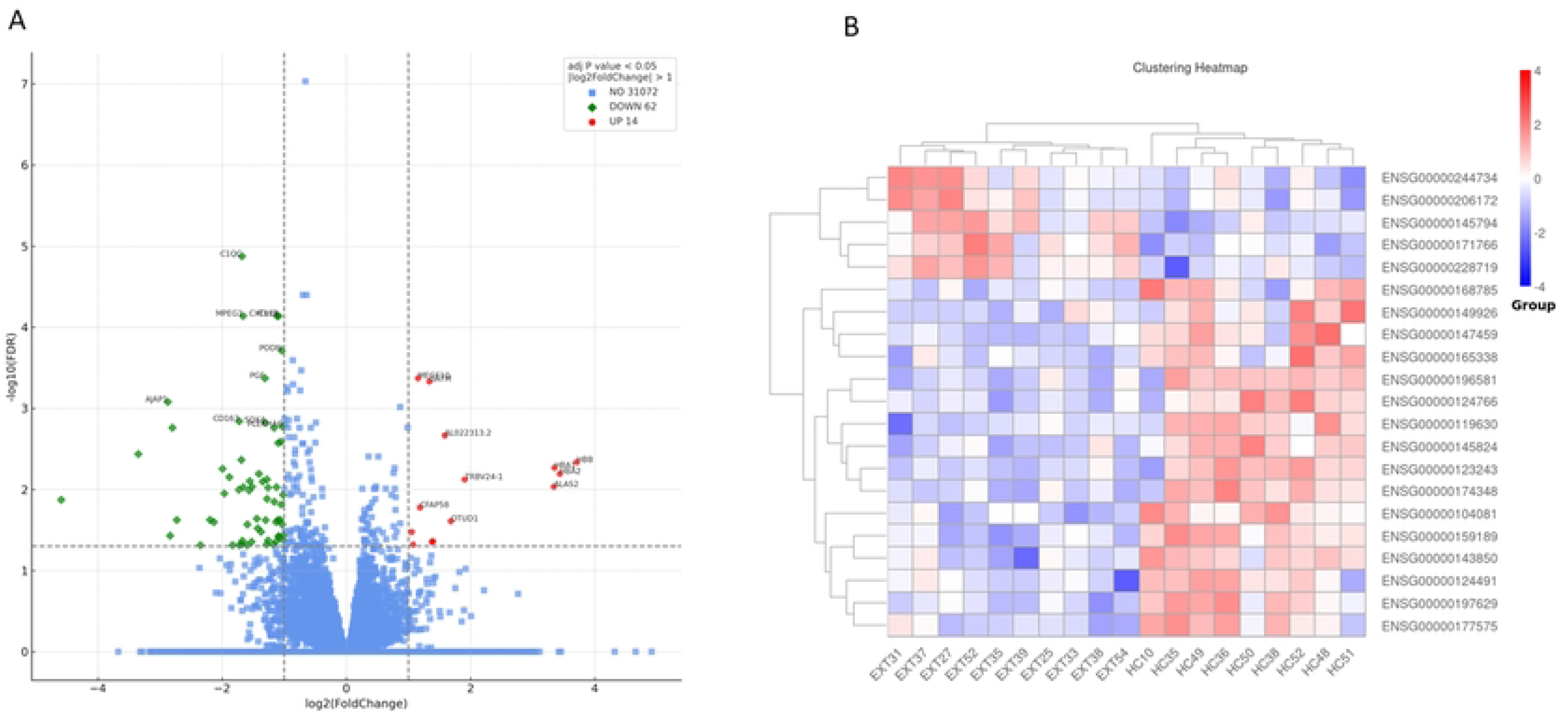
Differential gene expression between CKD and healthy control skeletal muscle. **(A)** Volcano plot showing differentially expressed genes (DEGs) between CKD and control muscle based on adjusted p-value < 0.05 and |log_2_ fold change| ≥ 1. Upregulated genes in CKD are shown in red (n = 14), downregulated genes in blue (n = 62), and non-significant genes in grey. Selected top-ranking DEGs by significance are annotated. **(B)** Heatmap of the top 20 DEGs ranked by adjusted p-value. Z-score normalised expression values are shown across all samples (CKD, n = 10; controls, n = 9), with unsupervised hierarchical clustering applied by row. Red indicates higher relative expression and blue indicates lower relative expression.

### Differential gene expression analysis

To gain initial insight into potential functional mechanisms, we examined the top 20 DEG’s (up or down regulated), ranked by adjusted p-value (Table 2). Several of these genes, including AJAP1, C1QC, DOCK5, MPEG1, ITIH5, TSPAN5, PGF, FAM57B and BMF, have no well-characterised role in skeletal muscle biology. Others, however, have known or plausible involvement in processes relevant to muscle health and pathology. For example, CXCL14^16^, PODN^17^, SOX4^18^ and MEGF10^19^ are linked to cell proliferation and differentiation, repair and regeneration; GATM^20^ is involved in creatine biosynthesis; HBB and HBA1 are components of haemoglobin^21^; and F13A1 has been documented to have a role in insulin sensitivity^22^. These top-ranking DEGs highlight a combination of known and potentially novel contributors to CKD-associated muscle wasting. Notably, unsupervised clustering of these genes revealed a clear separation between CKD and control groups in the heatmap (Figure 2B), supporting a distinct transcriptomic profile in CKD muscle.

**Table 2.**
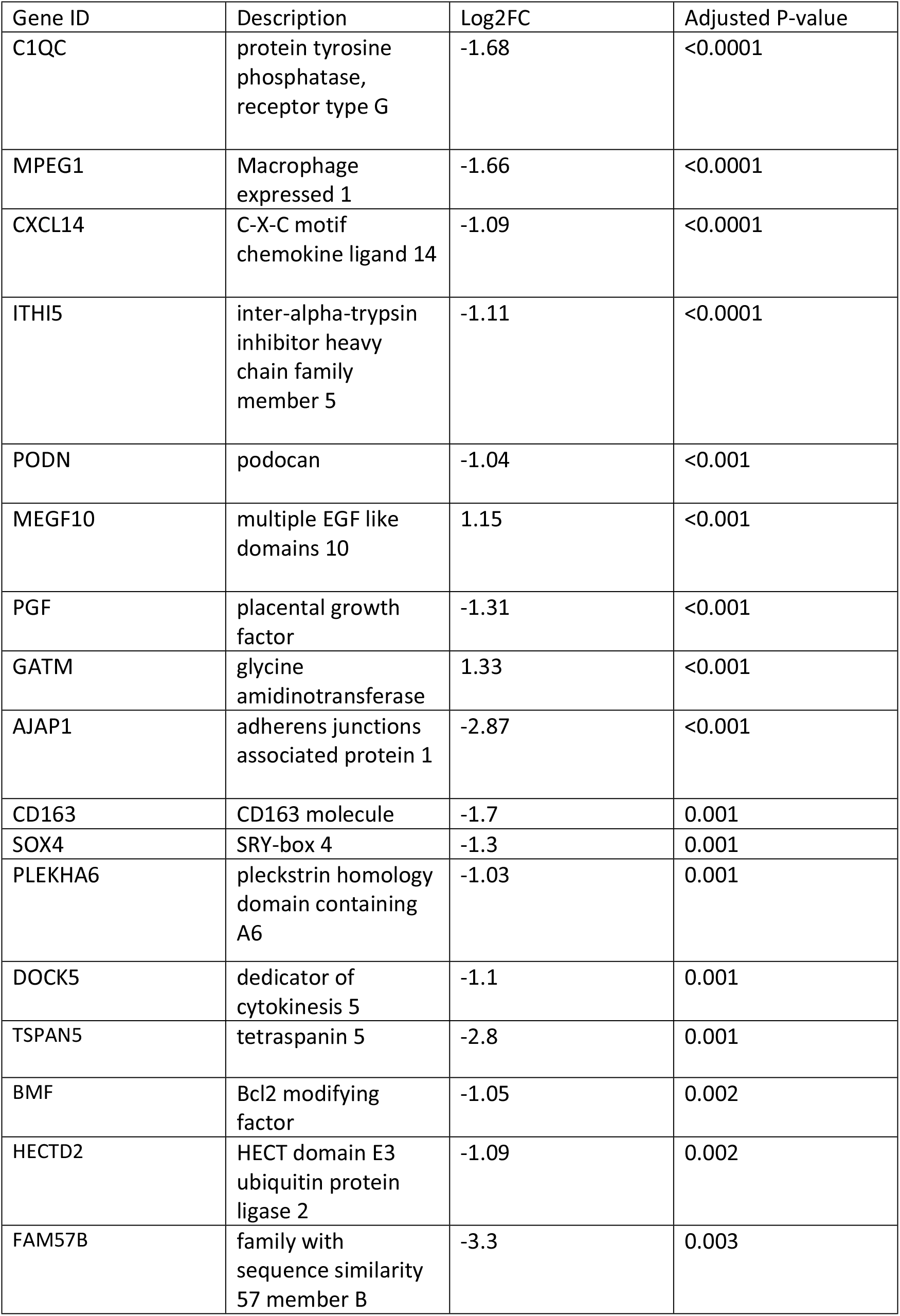
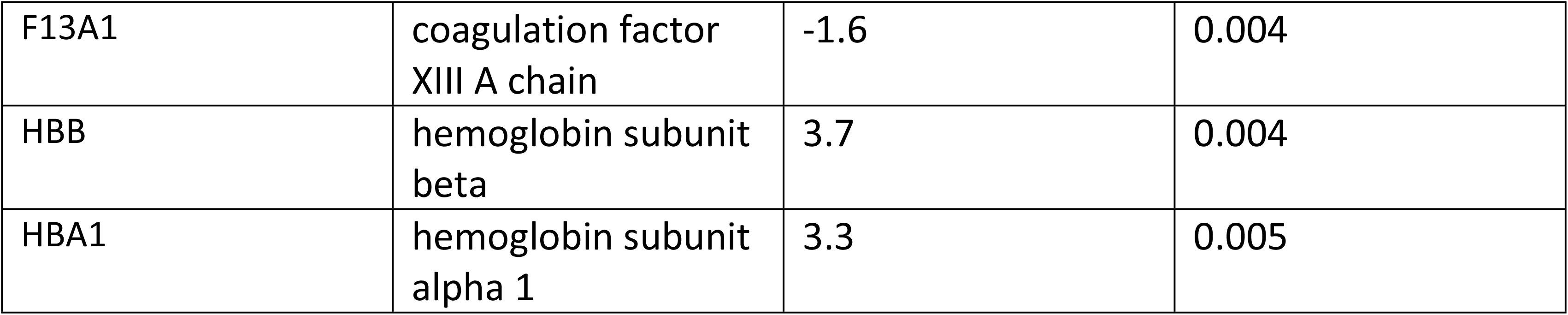
20 most differentially expressed genes, up or down-regulated, based on P values together with logFC and adjusted P values. The comparison is healthy controls to the CKD cohort, negative fold changes indicate downregulation in CKD and positive values indicate up regulation in CKD.

| Gene ID | Description | Log2FC | Adjusted P-value |
| --- | --- | --- | --- |
| C1QC | protein tyrosine phosphatase, receptor type G | -1.68 | <0.0001 |
| MPEG1 | Macrophage expressed 1 | -1.66 | <0.0001 |
| CXCL14 | C-X-C motif chemokine ligand 14 | -1.09 | <0.0001 |
| ITHI5 | inter-alpha-trypsin inhibitor heavy chain family member 5 | -1.11 | <0.0001 |
| PODN | podocan | -1.04 | <0.001 |
| MEGF10 | multiple EGF like domains 10 | 1.15 | <0.001 |
| PGF | placental growth factor | -1.31 | <0.001 |
| GATM | glycine amidinotransferase | 1.33 | <0.001 |
| AJAP1 | adherens junctions associated protein 1 | -2.87 | <0.001 |
| CD163 | CD163 molecule | -1.7 | 0.001 |
| SOX4 | SRY-box 4 | -1.3 | 0.001 |
| PLEKHA6 | pleckstrin homology domain containing A6 | -1.03 | 0.001 |
| DOCK5 | dedicator of cytokinesis 5 | -1.1 | 0.001 |
| TSPAN5 | tetraspanin 5 | -2.8 | 0.001 |
| BMF | Bcl2 modifying factor | -1.05 | 0.002 |
| HECTD2 | HECT domain E3 ubiquitin protein ligase 2 | -1.09 | 0.002 |
| FAM57B | family with sequence similarity 57 member B | -3.3 | 0.003 |
| F13A1 | coagulation factor<br>XIII A chain | -1.6 | 0.004 |
| HBB | hemoglobin subunit<br>beta | 3.7 | 0.004 |
| HBA1 | hemoglobin subunit<br>alpha 1 | 3.3 | 0.005 |

### Suppressed immune-related signaling in CKD muscle and an upregulation of oxygen transport capacity

To understand the function implications of all DEGs we carried out GO pathway enrichment and KEGG analyses on the 76 DEGs to further explore the biological processes dysregulated in CKD skeletal muscle. GO analysis showed several immune and inflammatory processes were downregulated, including regulation of leukocyte chemotaxis, mast cell degranulation, regulation of leukocyte migration and platelet activation. These were driven by downregulated genes such as CXCL14, C1QC, CCL2, and CD163, pointing to a possible attenuation of local immune cell signalling and recruitment (Figure 3A).

**Figure 3.**
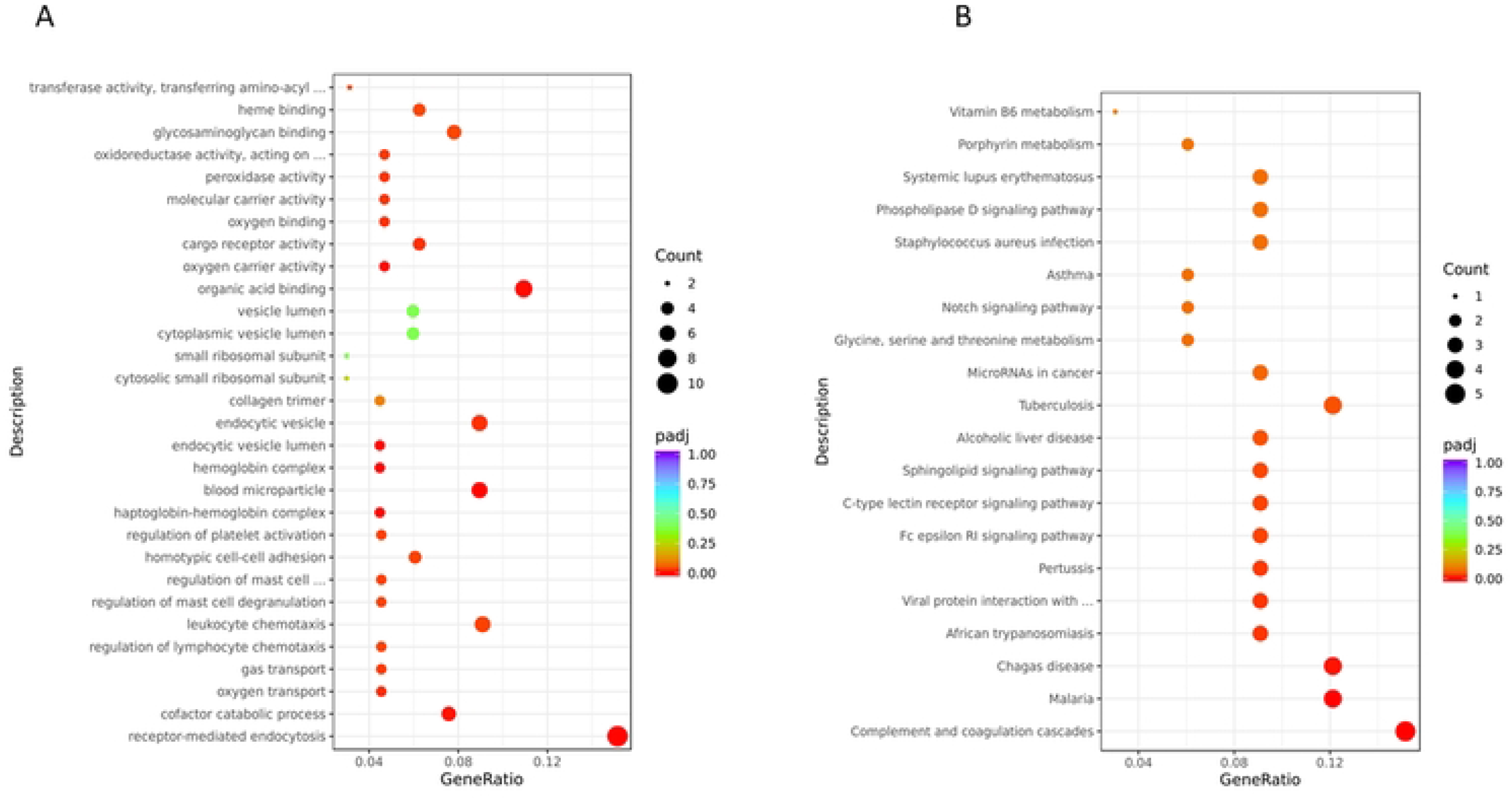
Functional enrichment analysis of differentially expressed genes in CKD skeletal muscle. **(A)** Gene Ontology (GO) enrichment analysis of 76 DEGs identified in CKD versus healthy control muscle (adjusted p < 0.05, |log_2_ fold change| ≥ 1). **(B)** KEGG pathway enrichment analysis of 76 DEGs identified in CKD versus healthy control muscle (adjusted p < 0.05, |log_2_ fold change| ≥ 1).

In support of this, KEGG analysis revealed several significantly enriched pathways, many of which align with the immune-related signals observed in the GO results. These included systemic lupus erythematosus, Staphylococcus aureus infection, and tuberculosis, pathways, commonly enriched in transcriptomic analyses due to the presence of genes related to immune surveillance, complement activation, and haemoglobin metabolism. These were primarily driven by downregulation of HBB, HBA1, CXCL14, C1QA, C1CB and CD163, reinforcing the finding of attenuated immune and redox signalling in CKD muscle (Figure 3B).

### Evidence of impaired regeneration and repair signatures

Although global enrichment analysis did not strongly highlight classical regeneration or muscle repair pathways, closer inspection of the top-ranking DEGs (Table 2) revealed transcriptional changes in several genes with known or plausible roles in these processes. For example, MEGF10, a gene associated with satellite cell function and muscle regeneration, was among the most significantly downregulated genes in CKD muscle. Similarly, the expression of CD163, a macrophage scavenger receptor involved in tissue repair and resolution was reduced. These changes support the notion of potential impairment of regenerative signalling in CKD muscle. However, there was no difference in expression of canonical proteins involved in skeletal muscle reparative pathways such as MyoD and Myogenin.

Taken together, these findings indicate that CKD may subtly impair key components of the muscle’s intrinsic regenerative and metabolic capacity. Although not dominant at the pathway level, these gene-level changes may contribute to the progressive decline in muscle quality and resilience observed in individuals with chronic kidney disease.

## Discussion

This study reports a distinct transcriptomic profile of skeletal muscle in patients CKD, revealing significant downregulation of immune and tissue repair pathways, alongside a small number of potentially compensatory upregulated genes.

One of the most prominent themes emerging from our data is the downregulation of immune-related genes and pathways in CKD skeletal muscle. Gene Ontology and KEGG enrichment analyses highlighted significant suppression of biological processes associated with leukocyte chemotaxis, cytokine receptor activity, and platelet activation, alongside enrichment of immune-related pathways such as systemic lupus erythematosus and Staphylococcus aureus infection, which reflect broad transcriptional downregulation of innate immune genes that also serve reparative or homeostatic functions.

Among the differentially expressed genes, CD163, an anti-inflammatory scavenger receptor expressed on M2-like macrophages and typically associated with tissue repair, was significantly downregulated in CKD. This contrasts with its well-established upregulation during chronic inflammation^23^ and with our previously published PCR-based data from the same cohort, which showed increased expression of IL-6, TNF-α, and CCL2, suggesting a state of intramuscular inflammation^8^. In the present RNA-seq dataset, however, these cytokines were either unchanged or downregulated, suggesting that overt inflammation may not be a consistent or dominant feature of CKD muscle. This discrepancy may reflect methodological differences (e.g., bulk RNA-sequencing vs. targeted PCR), or context-specific factors such as disease stage, metabolic status, or patient heterogeneity.

A reduced expression of CD163 may also reflect a reduced presence of resident macrophages or a blunted repair response in the CKD muscle environment. Supporting this, other immune-associated genes, including MPEG1 (macrophage-expressed gene 1) and CXCL14 (a chemokine involved in immune cell recruitment and regeneration), were also significantly downregulated. The downregulation of DOCK5 and DOCK10, which regulate leukocyte signalling and migration^24^, further reinforces a general attenuation of immune responsiveness. This attenuated immune transcriptional profile may suggest that CKD muscle may exist in a transcriptionally suppressed or immunologically quiescent state, impairing the muscle’s capacity to resolve damage and support regeneration in the CKD environment.

In addition to the downregulation of immune-related genes and pathways that may impair regenerative signalling, our data also provide further support for impaired muscle regeneration and repair capacity in CKD. Podocan, which has been shown to have a role in muscle regeneration^25^ was down regulated in CKD, together with SOX4, that has been shown to facilitate muscle myoblast differentiation^18^. Many of the DEGs with known roles in myogenesis and satellite cell activation were either downregulated or unchanged in CKD muscle compared to healthy controls. Key canonical markers of satellite cell activation and differentiation, including PAX7, MYOD1, *and* MYOGENIN, were not differentially expressed, together with down regulation of PODN, a repair-associated gene.

Interestingly, MEGF10, which plays a critical role in satellite cell-mediated muscle regeneration^19^, was significantly upregulated in CKD muscle. MEGF10 knockout mice have reduced muscle mass and neuromuscular junction instability^26^ suggesting an important role in maintenance of muscle health. This may represent a compensatory mechanism to preserve the regenerative potential of muscle in response to chronic muscle stress. We also reported downregulation of CXCL14. A recent study has shown that reduced CXCL14 expression may facilitate satellite cell activation and promote regeneration, especially in injury contexts^16^. Thus, the suppressed expression of *CXCL14* in CKD muscle may reflect an adaptive, pro-myogenic shift.

This regenerative profile likely reflects the resting, non-injured state of the individuals at the time the muscle biopsies were collected. This profile is difficult to interpret with lack of a stimulus that would stimulate these pathways. Our previous data has suggested that in an exercise naïve individual combined resistance and aerobic exercise is unable to activate processes of myogenesis, a process that was restored following training (Dougs paper), suggesting impaired regenerative responses. Interestingly, we did not detect any change in expression of key drivers of protein degradation such as myostatin, MuRF-1 or MAFbx, in line with our previous report^8^, suggesting there was no overt activation of skeletal muscle protein degradation under basal conditions.

A limitation of this study is the small cohort size, which may reduce the statistical power to detect genes influenced by CKD by small fold changes. To maintain a conservative approach, we applied a stringent threshold (adjusted p-value < 0.05 and |log_2_FC| ≥ 1) to define differentially expressed genes. While this enhances specificity, it may also exclude genes with more modest changes that contribute to disease pathology, particularly in a complex and multifactorial condition like CKD. the use of bulk RNA sequencing does not resolve gene expression at the level of specific cell types. Given the cellular heterogeneity of skeletal muscle—including myofibres, immune cells, endothelial cells, and fibro-adipogenic progenitors—some transcriptional changes may reflect shifts in cell composition rather than regulation within a specific cell type. Future studies incorporating single-cell or spatial transcriptomics could help disentangle these effects. Finally, transcriptomic data alone cannot provide insight into post-transcriptional regulation or protein-level effects. Some differentially expressed genes may not translate into functional changes, and integration with proteomics or histological data would be valuable to corroborate key findings.

In conclusion, we identified a distinct gene expression signature in CKD muscle characterised by suppression of immune and regenerative pathways, alongside potential compensatory upregulation of genes involved in oxygen transport. Notably, classical inflammatory mediators and myogenic regulators were either unchanged or downregulated, suggesting that CKD muscle may exist in a blunted or unresponsive state rather than a persistently inflamed one. These findings refine our understanding of the molecular landscape in CKD-associated muscle dysfunction and highlight potential targets for future mechanistic and therapeutic exploration. Larger, longitudinal, and functionally validated studies will be required to further elucidate the drivers and reversibility of these transcriptional alterations.

## Acknowledgements

This is a summary of independent research funded by the Leicester Hospitals Charity Kidney Care Appeal and the Stoneygate Trust, and supported by the National Institute for Health and Care Research (NIHR) Leicester Biomedical Research Centre (BRC). The views expressed are those of the author(s) and not necessarily those of the Kidney Care Appeal, the Stoneygate Trust, the NIHR or the Department of Health and Social Care. We thank Dr Douglas Gould and Dr Soteris Xenophontos for their help with the collection of the muscle biopsy samples. For the purpose of open access, the author has applied a Creative Commons Attribution license (CC BY) to any Author Accepted Manuscript version arising from this submission. There are no conflicts of interest to disclose.

